# Seminal Fluid Adipokinetic Hormone Increases Insemination Refractoriness in Female *Aedes aegypti*

**DOI:** 10.64898/2025.12.27.696674

**Authors:** Laura Sirot, Ferdinand Nanfack Minkeu, William Reid, Gemma Briggs, Dhwani Parsana, Meghan Wright, Jade Baek, Anna McGlade

## Abstract

Mating often changes behavior and physiology of female insects. In many species, these changes have been attributed to receipt of seminal fluid molecules (SFMs). SFMs influence phenotypes including feeding, egg production, and response to male courtship and insemination attempts. These same phenotypes are potential targets for management of insect pests. *Aedes aegypti* mosquitoes are the primary vector of several pathogens including dengue, Zika, and chikungunya. Within an hour after an initial insemination, female *Ae. aegypti* are generally refractory to subsequent inseminations, a response attributed to SFMs. However, the specific molecules involved in inducing long-term insemination refractoriness have not been identified. In a previous study, we identified adipokinetic hormone (AKH) precursor protein as an SFM in *Ae. albopictus.* AKH is a well-studied insect neuropeptide that impacts phenotypes including those related to metabolism, locomotion, and reproduction. In this study, we investigated whether AKH is an SFM in *Ae. aegypti* and whether it impacts female re-insemination patterns. We first established that AKH is produced in the male reproductive tract and transferred to females during mating, and is, therefore, an SFM. We then created an AKH-null line which allowed us to demonstrate that seminal fluid AKH contributes to long-term insemination refractoriness of females. Together, these results have established a novel expression pattern for AKH and identified AKH as a contributor to *Ae. aegypti* insemination refractoriness, laying the groundwork for understanding the evolution and mode of action of novel seminal fluid proteins as well as for investigating novel pathways or approaches for mosquito control.

## 1. Introduction

*Aedes aegypti* mosquitoes threaten much of the world’s population through transmission of pathogens including Zika, dengue, chikungunya, and yellow fever viruses (Achee et al., 2019; Kraemer et al., 2015; Shepard et al., 2011; Weaver, 2018). This species has adapted to thrive in urban zones, preferentially feeding on human blood and laying eggs above the water line in natural and artificial pools of standing water (Gubler, 2011; Harrington et al., 2001; Moyes et al., 2017; Scott et al., 1997; Weaver, 2018). Virus transmission occurs when infectious females feed on uninfected host blood (Clements, 2000). Although males do not directly spread pathogens, their biology is still important to understand because some control methods depend on male reproductive success and because males impact both female reproductive output and feeding behavior. After mating, females undergo transcriptional, behavioral, and physiological changes that modulate their vector capacity (Alfonso-Parra et al., 2014; Avila et al., 2011; Klowden, 1999; Nanfack-Minkeu et al., 2024; Nanfack-Minkeu & Sirot, 2022; Oliva et al., 2014). Relative to unmated females, mated females show changes including increased egg production, decreased blood-feeding and host-seeking behaviors, lengthened lifespan, and refractoriness to re-insemination (Gillott, 2003; Klowden, 1999; Kodrik, 2008; Lee & Klowden, 1999; Ramalingam, 1983; Villarreal et al., 2018). Therefore, understanding how males modulate female behavior is crucial for developing a successful population control strategy.

The efficacy of some mosquito population control techniques is strongly affected by female insemination patterns. For example, the sterile male technique depends on wild females mating with and using the sperm of lab-reared sterile males rather than those of wild fertile males (Kittayapong et al., 2019; Knipling, 1955). Female *Ae. aegypti* generally receive only a single ejaculate throughout their lives and are refractory to further insemination attempts (hereafter “insemination refractoriness”; Degner & Harrington, 2016; Oliva et al., 2014). However, females may receive ejaculates from multiple males under certain conditions including: (i) when a second insemination attempt takes place shortly after the first; (ii) when her first mate has previously inseminated many other females; and/or (iii) when a female is exposed to many males in close temporal sequence (Boyer et al., 2012; Dapples et al., 1974; Degner & Harrington, 2016; Dickinson & Klowden, 1997; Gwadz et al., 1971; Helinski et al., 2012; Williams & Berger, 1980; Young & Downe, 1982). Therefore, it is important to understand the mechanisms underlying variation between females in insemination patterns.

In *Ae. aegypti,* female insemination patterns, together with other post-mating responses, are modulated by seminal fluid molecules (SFMs) secreted from the male reproductive accessory glands (MAGs; Fuchs et al., 1968; Gwadz et al., 1971; Helinski et al., 2012; Klowden, 1999). MAGs impact insemination refractoriness in two phases: a short-term phase activated seconds post mating and a long-term phase which can last for a female’s entire lifespan (Craig, 1967; Fuchs et al., 1968; Helinski et al., 2012; Lee & Klowden, 1999; Villarreal et al., 2018). In the short term, gelatinous MAG secretions coagulate in the female reproductive tract, potentially blocking intromission, and the SFM Head Peptide I contributes within 5 minutes of the first mating to reduce re-insemination by about 20% (Duvall et al., 2017; Oliva et al., 2013). Specific SFMs contributing to long-term insemination refractoriness have not yet been identified. However, previous studies have identified protein fractions of approximately 7, 30, and 60kD that contribute to this long-term phenotype (Fuchs et al., 1969; Fuchs & Hiss, 1970; Gillott, 2003; Hiss & Fuchs, 1972; Lee & Klowden, 1999). These results suggest that long-term insemination refractoriness in female *Ae. aegypti* may be induced by a combination of partially redundant SFMs.

Our previous work, together with the work of other researchers, identified 280 SFPs in *Ae. aegypti* and 198 SFPs in *Ae. albopictus,* respectively (Boes et al., 2014; Degner et al., 2019; Sirot et al., 2011). Of these proteins, one candidate for regulating female insemination refractoriness is the adipokinetic hormone (AKH) (Boes et al., 2014). AKH is an 8-10 amino acid neuropeptide produced by many insects as well as other invertebrates, and is generally expressed in the *corpora cardiaca* (Gäde & Auerswald, 2003). The AKH peptide is a cleavage product of a larger precursor protein. Unexpectedly, in *Ae. albopictus,* the C-term portion of the precursor protein is transferred to females during mating (Boes et al., 2014). The AKH peptide itself has not yet been detected as transferred to females using mass spectrometry, but this could be due to its small size and post-translational modifications (Degner et al., 2019). AKH (AAEL011996) mRNA is highly expressed in the male abdomen of *Ae. aegypti* (Kaufmann et al., 2009). Further, the predicted *Ae. aegypti* AKH precursor protein is 6.4 kD (8.8 kD including its signal peptide) and thus is a candidate for the bioactive 7.6 kD SFM identified by Lee and Klowden (1999).

Although AKHs influence a variety of physiological processes (Kodrik, 2008), they are best known for their roles in regulating carbohydrate and lipid levels in the hemolymph through metabolism and/or release from the fat body during periods of stress (Gäde & Auerswald, 2003). In the mosquitoes studied to date (*Aedes aegypti* and *Anopheles gambiae*), females injected with AKH had increased hemolymph carbohydrate levels and decreased glycogen levels (Afifi et al., 2023; Kaufmann & Brown, 2008). In *An. gambiae*, AKH injection also increased female flight performance. In *Culex pipiens pallens*, knockdown of the AKH receptor (AKHR) in females resulted in higher incidence of abnormal ovarian morphology, decreased yolk and iron levels in the ovaries, and reduced numbers of follicles and laid eggs (Wang et al., 2025). AKH also impacts reproductive function in other groups including controlling male courtship and female fecundity in *Bactrocera dorsalis* (Hou et al., 2017), inhibiting egg maturation in *Gryllus bimaculatus* (Lorenz, 2003), modulating egg laying in *Caenorhabditis elegans* (Lindemans et al., 2009), and regulating male courtship and female pheromone production in a nutrition-dependent manner in *Drosophila melanogaster* (Lebreton et al., 2016).

In addition to these reproduction-related phenotypes, AKH/AKHR’s roles in reproduction are also suggested by their similarities to gonadotropin-releasing hormone (GnRH) and its receptor. GnRH and its receptor are key regulators of vertebrate reproductive processes (Flanagan & Manilall, 2017). Sequence alignments, evolutionary reconstructions, and receptor activation studies suggest that AKHR is the invertebrate homolog of the gonadotropin-releasing hormone (GnRH) receptor and that AKH and GnRH arose from a common ancestor(Lindemans et al., 2009, 2011; Plachetzki et al., 2016; Staubli et al., 2002). *Aedes aegypti* AKH receptors (AKHR-I and AKHR-II) are GPCRs with expression in many parts of the body including reproductive tissues and are activated by the AKH peptide (Afifi et al., 2023; Kaufmann et al., 2009; Marchal et al., 2018; Oryan et al., 2018). Together, this evidence suggests that seminal fluid derived AKH (sfAKH) could contribute to post-mating responses in female *Ae. aegypti*.

In the current study, we tested the hypothesis that seminal fluid AKH contributes to the insemination refractoriness of female *Ae. aegypti*. We first tested for expression of the AKH precursor protein in the male *Ae. aegypti* reproductive tract and transfer of the AKH peptide to females during mating. We then tested for the impact of sfAKH on insemination and remating behavior of females.

## 2. Materials and Methods

### 2.1 Overview

We used HPLC-MS to test whether the AKH peptide is transferred from males to females during mating. Once AKH peptide transfer was confirmed, we used an AKH-null strain created using CRISPR-Cas9 gene editing to test for impacts of male-derived AKH on female re-insemination. Since the AKH-null strain is knocked out for AKH not only in their MAGs but also in their *corpora cardiaca*, we isolated the impact of sfAKH in two ways: (i) we injected MAG homogenates from AKH-null and wildtype males into unmated females and tested for insemination and (ii) we injected AKH or a control peptide into unmated females and tested for insemination. After finding an effect of AKH on reducing insemination, we further investigated the detailed behavioral interactions between males and females previously mated to either AKH-null or wildtype males.

### 2.2 AKH-null Line Establishment

Four sgRNAs were synthesized *in vitro* and complexed to Cas9-NLS protein (PNABio, Thousand Oaks, CA) at equimolar concentrations to obtain an injection mix containing 300 ng/µL protein and 80 ng/µL of pooled sgRNAs (Kistler et al., 2015). The primers used in the study are listed in Table 1 and diagrammed in Figure 1. A total of 348 pre-blastoderm *Ae. aegypti* embryos (strain LVPIB12) were injected at the Insect Transformation Facility, University of Maryland and hatched five days following injection. A total of 65 surviving G0 were reciprocally backcrossed to wildtype LVPIB12 *Ae. aegypti* at a ratio of 1:2 and 1:10 for G0 females and males, respectively. Following mating, all backcrosses were pooled and the females were provided with a bloodmeal and allowed to oviposit individually in 30 mL Drosophila vials to obtain single-parent G1 populations. The G1 populations were hatched and reared individually to fourth instar, at which point a pooled sample 10 larvae were sampled from each of the populations and assessed for the presence of a CRISPR/Cas9-mediated deletion in AAEL011996 using PCR and Sanger sequencing trace decay analysis. Positive pools were sibling mated and females were allowed to oviposit individually in 30 mL Drosophila vials until a homozygous deletion line could be established. Genotyping of the mutant line was performed using a restriction fragment length polymorphism (RFLP) BstNI digest of the PCR product, where the mutant line was refractory to digestion due to loss of the BstNI site. Hereafter, this line is called the “AKH-null” line.

**Figure 1.**
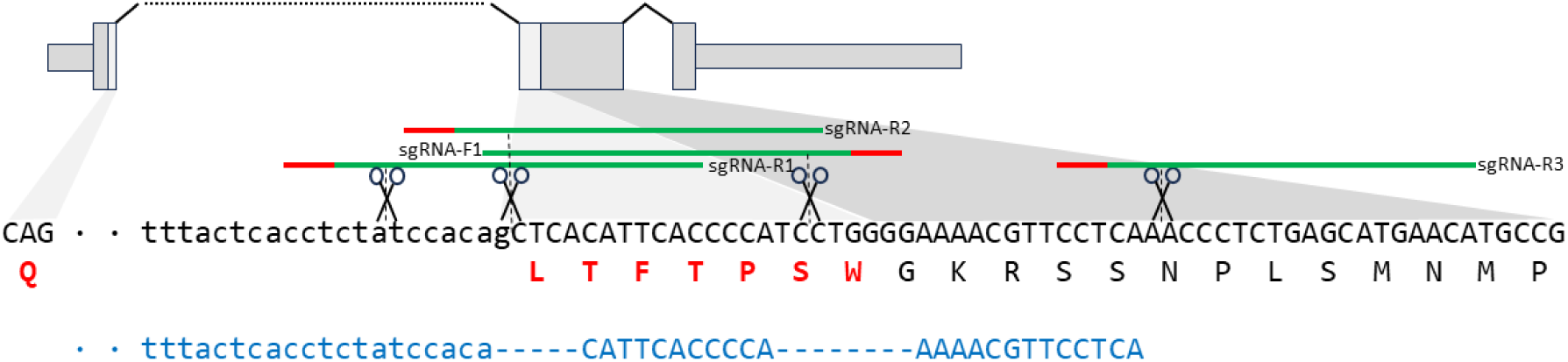
Schematic for the targeting of adipokinetic hormone (AKH; AAEL011996) for knock-out, and sequence of the deletion mutant. Wildtype sequence is written in black, mutant deletion sequence is written in blue. Amino acids highlighted in red indicate the AKH peptide of interest. Nucleotides indicated in lowercase represent intron I, green lines overlap the CRISPR/Cas9 sgRNA target sites with their respective PAM sequences indicated in red. The positionings of the scissors indicate the expected cleavage site of Cas9.

### 2.3 Mosquito Rearing

*Aedes aegypti* mosquitoes of the wildtype Liverpool (WT), AKH-null, dsRed, and Thai lines were raised in a colony at 27°C, >50% relative humidity and a 12h light: 12h dark photoperiod. dsRed males have the fluorescent dsRed peptide sequence fused to the β2 tubulin promoter, a sex-specific protein essential for spermatogenesis that localizes to the testes and results in sperm that fluoresce at 561 nm ( provided by L. Harrington lab; Smith et al., 2007). The Thai line was obtained from the Harrington lab (Cornell University) in 2019 and was generated from wild-collected egg (collected by Alongkot Ponlawat from Nakhon Ratchasima Province, approximately 260 km northeast of Bangkok). For all experiments, eggs were hatched in low dissolved oxygen water and larvae were reared in 1L of reverse osmosis water with 4 pellets of ground Hikari Cichlid Gold fish food. Pupae were separated into individual 5 mL tubes topped with cotton. Adults were released into sex specific 5L cages with screen tops and 10-12% sucrose water ( ≤ 100 individuals per cage).

**Table 1.**
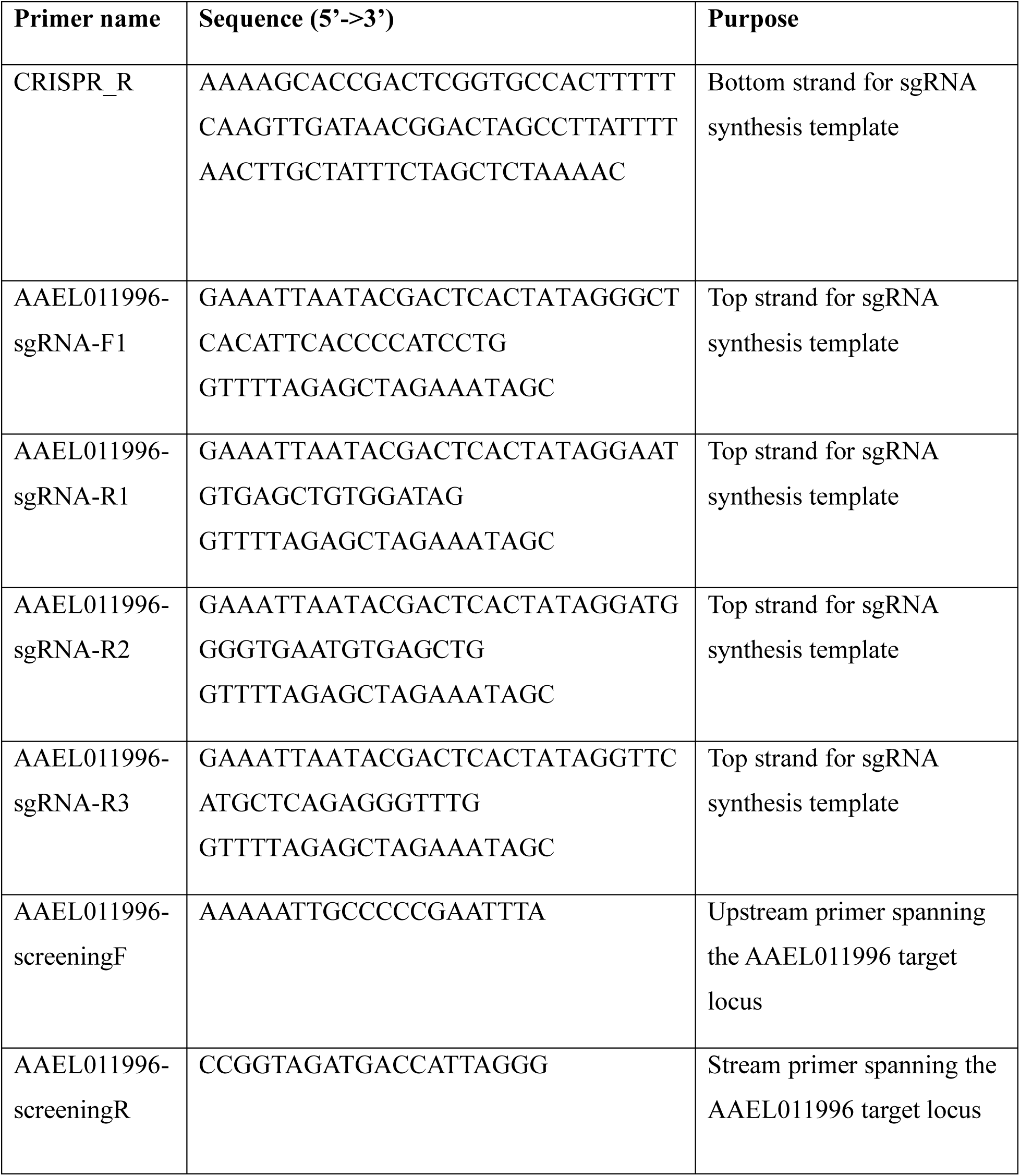
List of oligonucleotides used for the synthesis of the CRISPR/Cas9 sgRNAs and for genotyping of the mutant *Ae. aegypti* Adipokinetic hormone (AAEL011996) knock-out line.

### 2.4 Testing for the presence of the AKH peptide in male and female reproductive tracts

#### 2.4.1 RT-PCR

For whole mosquito RT-PCR, RNA was extracted from 8-9 two-week old male mosquitoes with legs and wings removed. For tissue-specific RT-PCR, tissues were dissected from unmated two-week old mosquitoes into 1x DEPC treated PBS. Total RNA was extracted using the RNeasy total RNA kit (Qiagen, Hilden, Germany) according to the manufacturer’s protocol. RNA was eluted in 30 uL RNase-free water and quantified using the NanoDrop 1000 where it was then stored at −80℃ until PCR. Reverse transcription and PCR of extracted RNA was performed using the OneStep RT-PCR kit (Qiagen, Hilden, Germany) according to the manufacturer’s protocol. Each 50 uL reaction contained 10 uL of 5X QIAGEN OneStep RT-PCR Buffer (12.5 mM MgCl2), 400 uM dNTP, 0.6 uM of either the AKH or the RPS7 forward and reverse primers (Table 2), 2 uL QIAGEN OneStep RT-PCR Enzyme Mix, 30-60 ng of extracted RNA, and RNase-free H_2_O. The RPS7 primers were used as an internal control to ensure the integrity of the RNA and a negative control was also prepared which substituted RNase-free H_2_O for RNA. The thermocycle conditions were as follows: Reverse transcription: 30 minutes at 50C, PCR: 15min at 95C, 30 cycles of 30 seconds at 94C, 30 seconds at 50C, and 1 minute at 72C, and then 10 minutes at 72C. Samples were run in a 2% agarose gel and visualized using GelRed and a BioRad Imager.

**Table 2:**
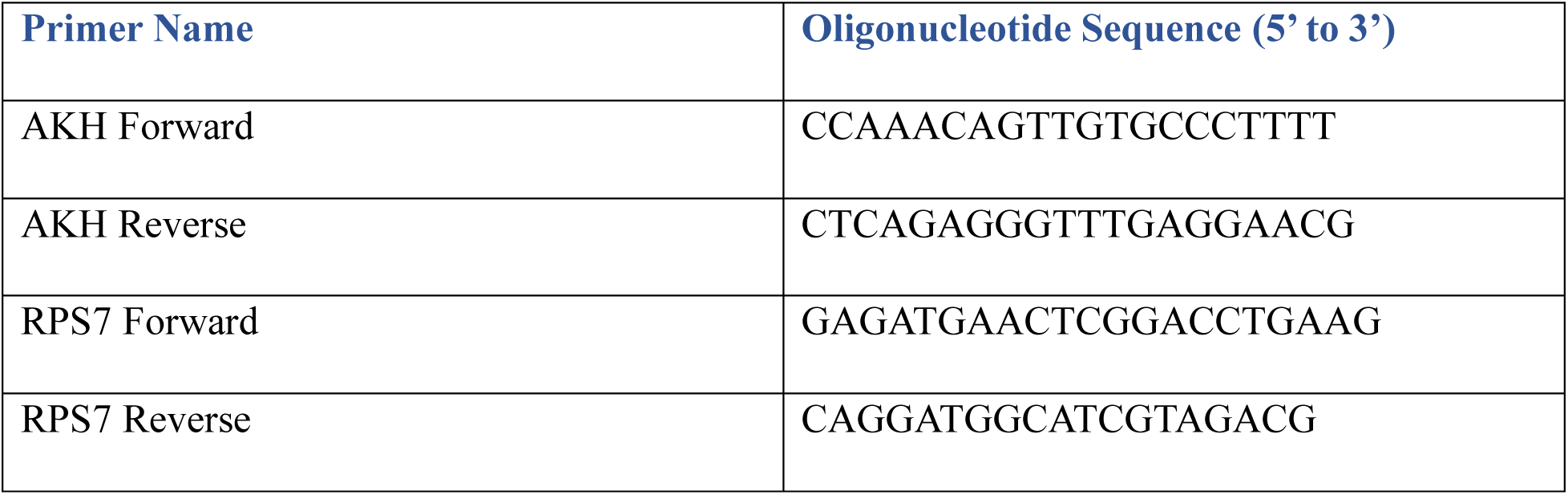
Primers used for reverse transcription PCR to examine gene expression of adipokinetic hormone.

#### 2.4.2 Western Blotting

Male accessory glands (MAGs) were dissected from 50 unmated 11-day-old WT or AKH-null males per sample. Dissections were conducted on ice in 1xPBS. MAGs were ground in 20µL RIPA lysis buffer with protease inhibitors (cOmplete^TM^ Protease Inhibitor Cocktail; Roche, Indianapolis, IN, USA). Twenty microliters of 2x Sample Buffer were added to each sample. The sample was boiled for 4 minutes.

The samples were separated using PAGE-SDS in a Mini-Protean System (BioRad) and transferred to a PVDF membrane. The membrane was cut at 25kD so that we could probe for AKH on the lower size range and probe for tubulin as a loading control on the upper size range. We used polyclonal primary antibodies for the AKH peptide (1:250) and for tubulin (1:2000). We used HRP-conjugated secondary antibodies (1:5000; goat anti-rabbit for AKH and goat anti-mouse for tubulin). We then applied Clarity ECL (BioRad, California) and visualized the chemiluminescence using a BioRad ChemiDoc imager.

#### 2.4.3 HPLC-MS

Male accessory glands and seminal vesicles (MAG/SV) and female lower reproductive tracts (LRTs) were dissected from 5-7-day-old *Ae. aegypti* WT mosquitoes. The tissues were dissected from unmated and mated females and unmated males into ultrapure water treated with proteinase inhibitor (cOmplete^TM^ Protease Inhibitor Cocktail; Roche, Indianapolis, IN, USA) under a dissecting microscope (4X). Mating was described in Parsana et al. (2022). Two replicates of this experiment were conducted. The first replicate consisted of tissues from 200 males, 184 unmated females, and 170 mated females. The second replicate consisted of tissues from 198 males, 191 unmated females, and 160 mated females. They were crushed using pestles and centrifuged at 12 000 RPM at 4°C for 10 minutes. Following the centrifugation, the supernatant was transferred into new clean tubes and brought up to a volume to make a final concentration of 1 individual/µL. The supernatant was used for the quantification of AKH using high-performance liquid chromatography with tandem mass spectrometry (HPLC-MS). For a positive control, we used a 2.5ng/mL standard of synthetic AKH peptide dissolved in ultrapure water treated with a protease inhibitor cocktail as described above. For negative controls, we used ultrapure water treated with protease inhibitor cocktail as described above.

Mass spectrometry analysis was done using an Agilent 1200/6410 HPLC-MS/MS (QQQ). Separation was performed using an Agilent Poroshell 120 EC-C18 column (2.1 x 100 mm, 2.7 μm particle size). Mobile phases were 100% water with 0.1% formic acid and methanol with 0.1% formic acid. A 7 min binary gradient was started at 60% water 40% methanol ramped to 100% methanol with a flow rate of 0.25 mL/min and a column temperature of 40°C. Injection volume was 30 μL. The mass spectrometer was run in positive mode with +3000V ESI at 350°C drying temperature with MRM detection. Transitions for the peptide were 961.5 > 573.9 *m*/*z* (qual ion) and 961.5 > 388.2 *m*/*z* (quant ion) both using a 25 V collision energy and 4 V cell accelerator voltage.

### 2.5 Testing for effects of mating with AKH-null males on subsequent female re-insemination

Matings were performed in a low light environment at 25°C and ≥ 50% humidity, using 2-5 day old mosquitoes. All mosquitoes used were previously unmated and females were WT. Females were alternately assigned to one of three treatments: unmated, mated to WT males, or mated to AKH-null males. Females in the unmated treatment were aspirated into a new cage with 10% sucrose water. Females in mating treatments were individually aspirated into cages of 20 males and observed until a single complete mating of at least six seconds occurred (Spielman et al., 1967). Females that did not mate for at least six seconds were excluded from analysis. After mating, females were transferred to group cages according to treatment with 10% sucrose water. Approximately 24 hours later, a second mating opportunity (first for the unmated treatment) was presented to all three groups of females by adding 25-35 4-7 day old dsRed unmated males to each cage of 20 females. These mosquitoes were left to mate *ad libitum* for 24 hours. Males were then removed from the cage and the remaining females were frozen at −20 degrees before dissection.

Spermathecae and lower reproductive tract were dissected from females in 1x PBS and mounted on uncharged glass slides, using approximately 25µL ProLong Gold Antifade mountant with DAPI stain (ThermoFisher). Slides were sealed with 3 coats of clear varnish and dried for at least 24 hours. Samples were imaged using an Olympus Fluoview FV3000 Confocal Microscope. Images using a 20x objective were visualized using the following parameters: 405 nm (DAPI) (1%), 700 HV, 1.0x gain, 0% offset and 561 nm (dsRed) (1.6%), 650 HV, 1.0x gain, and 0% offset. Individuals were scored for presence of WT and dsRed-labeled sperm in their spermathecae. Three replicates of this experiment were conducted with a total of 50-58 females scored for each treatment. Chi squared tests were used to test for differences in the proportion of females that were re-inseminated in each of the treatment groups.

### 2.6 Testing for effects of injection with MAGs from AKH-null males on subsequent female insemination

#### 2.6.1 MAG Homogenate Preparation

Male accessory glands and seminal vesicles (MAG/SV) were dissected from 40 WT unmated males and 40 AKH-null unmated males that were 7 to 8 days in dissecting saline solution (0.13 M NaCl, 4.7 mM KCl, 1.9 mM CaCl_2_; Ephrussi & Beadle 1936). Immediately after dissection, the SV and MAG were ground with a pestle in 1.5 mL tubes containing 40µL of Aedes Saline. The tubes were sonicated using an ultrasonicater (Fisher Scientific, Pennsylvania) for 15 seconds. The samples were then centrifuged at 0°C for 15 minutes at 12,000 rpm. The supernatant was transferred to a fresh sterile tube. The volume was brought up to 40µL for each sample using Aedes Saline to get 1 male-equivalent/µL. Samples were stored at −20 °C for up to a month until ready to be injected.

#### 2.6.2 Injections

Four to five day old unmated WT females were used for the injections. The females were starved for 12 hours prior to the injections to avoid bursting of the female due to excessive liquid inside the body. The females were anesthetized by placing them in glass tubes on ice for ∼5 mins and maintaining them on ice during the injections. The females were injected through the thorax using a Nanoject II (Drummond Scientific Company, Pennsylvania) (Helinski et al., 2012). For MAG/SV injections, 40-50 females were injected per treatment per replicate with either WT MAG/SV, AKH-null MAG/SV, or Aedes Saline. The females were injected with the equivalent of ¼ of a male’s MAG/SV in 250nL. Injected females were put in recovery cages containing 10-12% sucrose water and the cages were covered with a damp cloth and plastic wrap to provide humid conditions. The females were left in the recovery cage for ∼24 hours before testing them for mating.

#### 2.6.3 Insemination Assay

Individual females from recovery cages were transferred to mesh-topped 0.5L ice cream carton cages which contained two unmated 6-day-old WT males. The cartons were labeled with numbers which had been randomly assigned using a random number generator so that the researcher would be unaware of the treatment group during scoring. The males and females were allowed to mate ad libitum for ∼48 hours. Cotton balls dipped in 10-12% sucrose water were placed on the top mesh and were refreshed every 24 hours. After ∼48 hours, the small individual cages were placed at −20 °C for ∼2 to 5 mins and females were dissected to obtain their spermatheca in Aedes Saline. The spermathecae were crushed using individual cover slips and sperm were visualized using a compound microscope at 10X and 40X magnification (Olympus, Pennsylvania). Chi squared tests were used to test for differences in the proportion of females that were inseminated in each of the treatment groups.

### 2.7 Testing for effects of injection with AKH peptide on subsequent female insemination

This experiment followed the same protocol as that of Section 2.6 except females were injected with synthetic peptides rather than MAG/SV. AKH peptide (sequence: {pGLU}LTFTPSWa) and a control scrambled peptide (sequence: {pGLU}TFSPTWLa) were used at two different concentrations, 8 picomoles/250 nL (8A and 8S) and 80 picomoles/250 nL (80A and 80S) dissolved in Aedes Saline for the peptide injections. The peptides were synthesized by GenScript. Injection and insemination assay procedures were as described above in section 2.6.2. This experiment was repeated using females from both the LVP and the Thai strain. For peptide injections, 29-40 females were injected per treatment per replicate.

### 2.8 Testing for effect of mating with AKH-null males on subsequent female interactions with males

Because we found effects of sfAKH receipt on subsequent insemination patterns, we next investigated whether these effects might be mediated by differences in mating behavior of females previously mated to males who did or did not transfer sfAKH. In this study, detailed observations were made of intersexual interactions involving females from one of three treatments: previously unmated, previously mated to WT males, or previously mated to AKH-null males. The methods for obtaining these treatment groups were the same as those described in Section 2.5 above. Twenty-four to 48 hours later, females were observed in interactions with WT males. For this step, the three treatments were labeled with codes so that the observer was unaware of the treatment group. Individual females were aspirated into a transparent cage covered with mesh containing three WT males. The order of female treatments used was randomized. Mating duration was recorded in seconds. Genital pairings between the female and one of the males that lasted at least six seconds were recorded as successful matings ((Spielman et al., 1967). The total duration of the trials was four minutes. The top of cage was tapped gently every 15 seconds during the trial to encourage the mosquitoes to interact with each other. A chi-square analysis was used to test for differences in proportion of females that mated. A Kruskall-Wallis test was used to test for differences in mating duration, followed by post-hoc Mann-Whitney U tests for pairwise comparisons.

## 3. Results

### 3.1 AKH-null line establishment

Sequencing of one of the mutant lines identified two deletion lesions of five and eight nucleotides, respectively positioned at the beginning of exon II and present in the AKH peptide coding region. In addition, the deletion disrupted the terminal G of intron I.

**Table 2:**
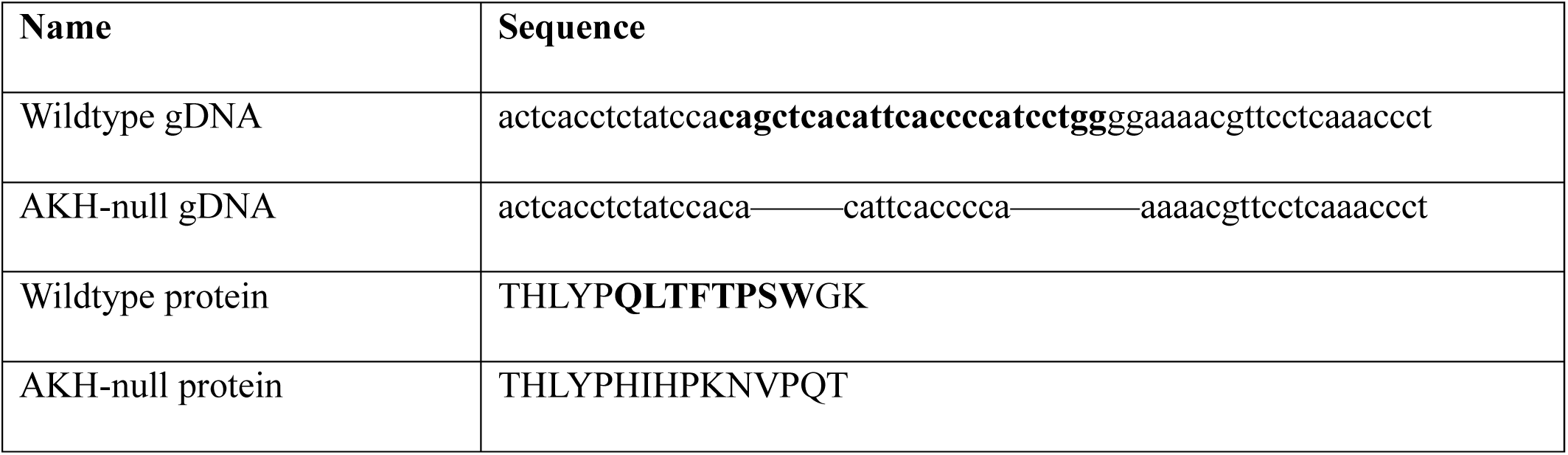
Adipokinetic hormone (AKH) protein and cDNA sequence of wildtype and CRISPR-generated AKH-null line. Bolded sequence encode the AKH peptide.

### 3.2 AKH is highly expressed in the MAG/SVs of wildtype males

#### 3.2.1 RT-PCR

We next tested for expression of the AKH gene in both the wildtype and CRISPR-Cas9 modified lines. RT-PCR analysis of tissues from wildtype mosquitoes showed high expression in the male accessory glands and seminal vesicles and moderate expression in the female head and thorax (Figure 2A). RT-PCR analysis of whole mosquitoes from the CRISPR-Cas9 modified line showed no AKH expression (Figure 2B). This line became our AKH-null line for the remaining experiments.

**Figure 2.**
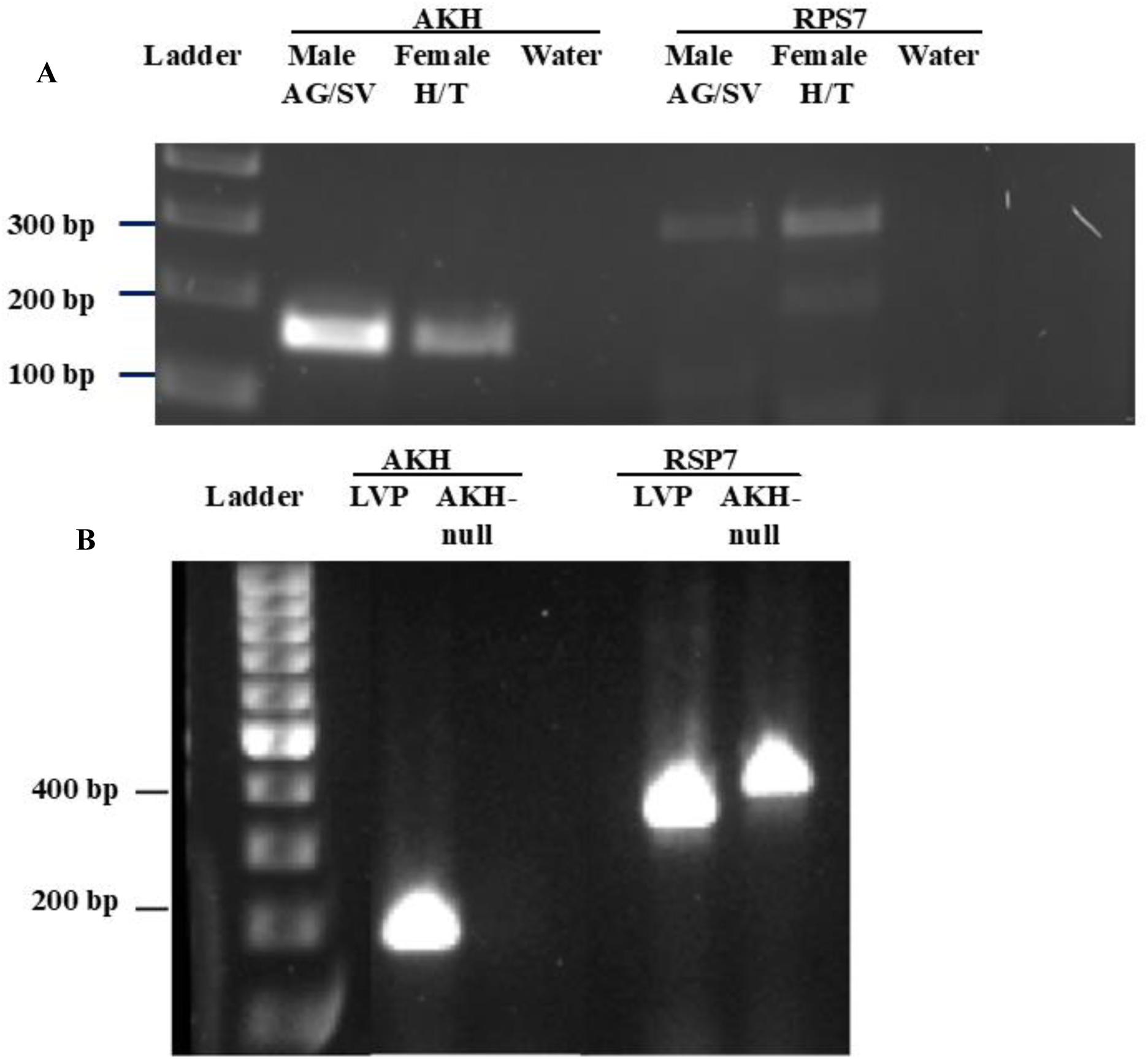
Reverse transcription PCR results: A. AKH and RPS7 transcript expression in the male accessory glands and seminal vesicles (MAG/SV) and female head and thorax (H/T) of Liverpool (wildtype) mosquitoes. B. AKH and RPS7 transcript expression in the whole bodies of males from the LVP strain (wildtype) and males from the AKH CRISPR deletion strain (AKH-null). RPS7 from the LVP and AKH-null lines ran at different sizes due to a difference how GelRed binds to the total amount of DNA present (https://biotium.com/wp-content/uploads/2018/11/PI-41011.pdf). Representative gels for three replicates of this assay.

#### 3.2.2 The AKH precursor protein is found in male MAG/SVs

Next, we tested whether the AKH protein was produced in the MAG/SVs of wildtype and AKH-null lines. Western blotting showed that the presence of a band at approximately 7.5kD was produced in the MAG/SVs of wildtype males but not of the AKH-null males (Figure 3). This molecular weight corresponds to the weight of the full AKH precursor protein before processing occurs.

**Figure 3.**
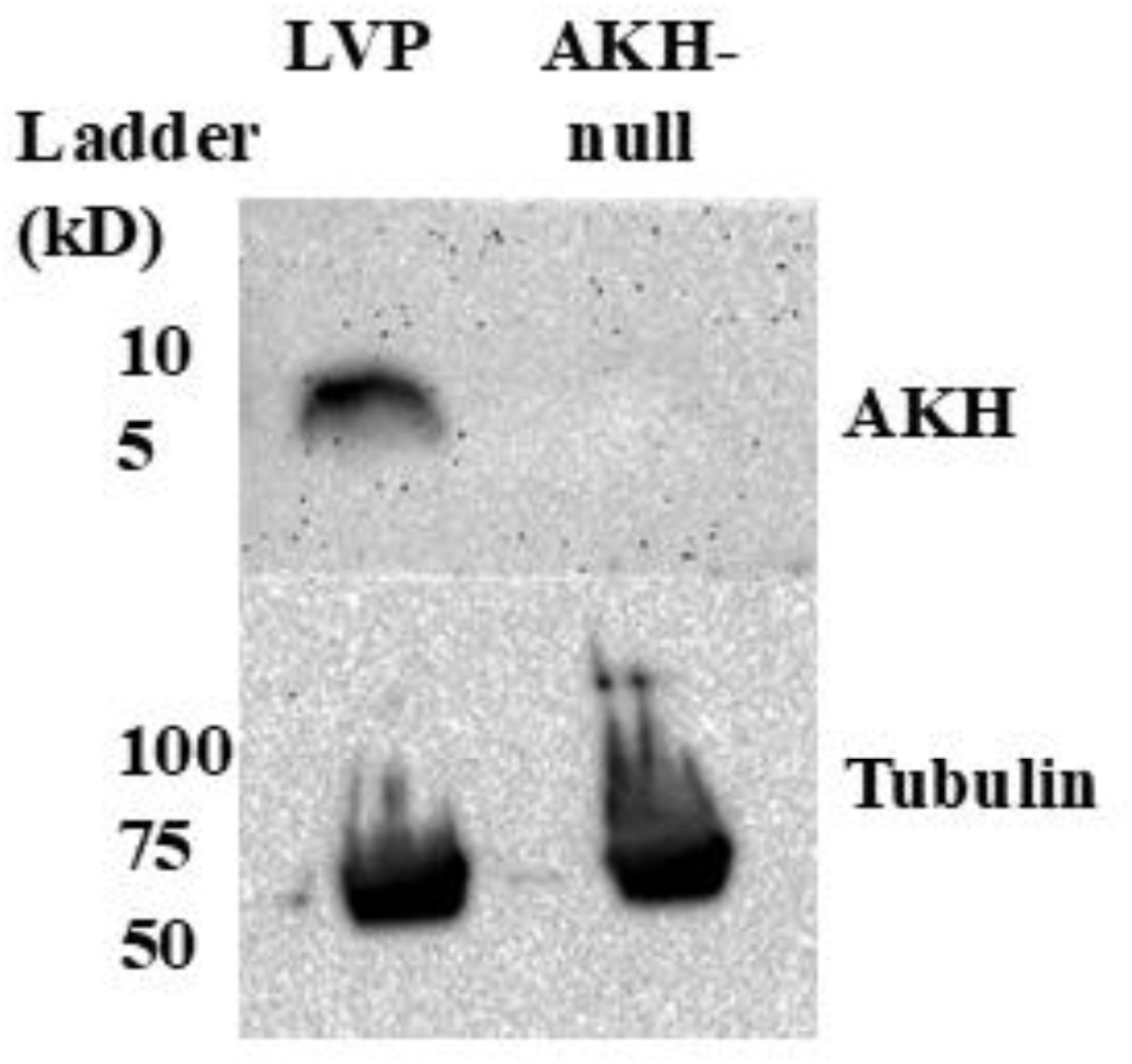
Western blot showing protein levels of AKH and tubulin (as a loading control) from the male accessory glands and seminal vesicles of LVP (wildtype) males and males from the AKH CRISPR deletion line. Samples were from 50 males each. Representative blot for three replicates of this assay.

#### 3.2.3 AKH peptide is transferred from males to females during mating

After establishing that AKH mRNA and protein is present in the MAG/SVs of wildtype males, we tested whether it is transferred to females during mating. Two independent replicates of the HPLC-MS analysis showed a distinct peak at 7.7-7.8 minutes in both the synthetic AKH (Figure 4A) and MAG/SV (Figure 4B) samples. The protease negative control (data not shown) and the unmated females (Figure 4C) samples showed no peak at this time. The mated female samples showed a small but distinct peak at 7.7-7.8 minutes (Figure 4D).

**Figure 4.**
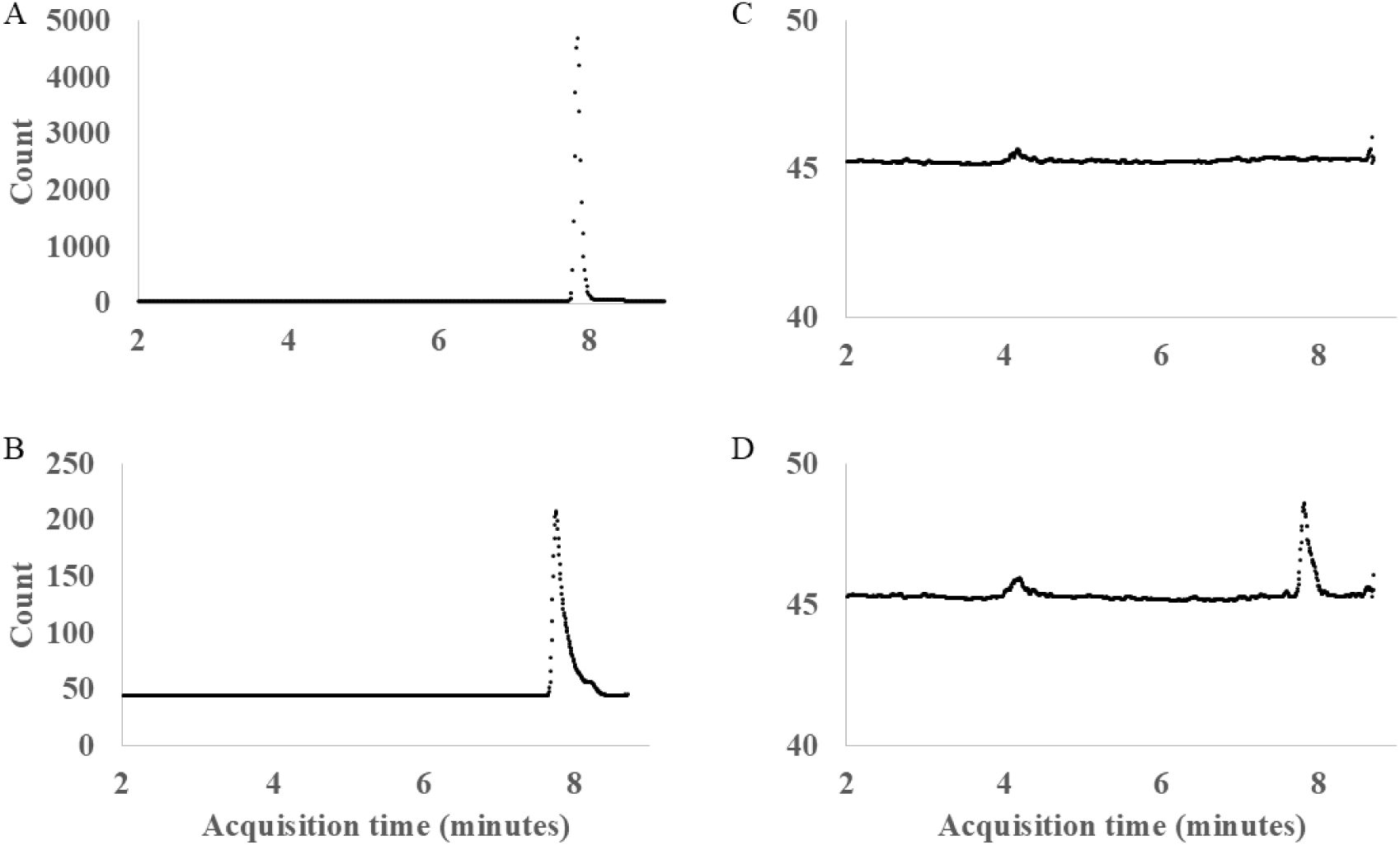
HPLC analysis of A: synthetic AKH peptide standard (2.5ng/mL); B: Wildtype (LVP) male reproductive accessory glands and seminal vesicles; C: Unmated female lower reproductive tracts; D: Mated female lower reproductive tracts.

### 3.3 Females previously mated with AKH-null males were more likely to be inseminated by a second male than females previously mated to wildtype males

Once it was established that AKH was transferred to females during mating, we tested whether females who received seminal fluid AKH (sfAKH) were less likely to be subsequently inseminated than females who did not receive sfAKH. In all three replicates of this experiment, 0-10% of females previously mated to WT males, 20-22% of females previously mated to AKH-null males, and 87-100% of previously unmated females were inseminated by a second male (Table 1). Insemination frequency did not differ between the replicates (*X^2^* = 0.64; *p* = 0.73), and, therefore, we combined the data from the three replicates for analysis. Females previously mated to AKH-null males were significantly more likely to be inseminated by a second male than females previously mated to WT males (*X^2^* = 5.01; *p* = 0.03). Females from both mated treatments also were significantly less likely to be inseminated by a second male than previously unmated females (WT vs unmated: *X^2^* = 78.4; *p* < 0.00001; AKH-null vs unmated: *X^2^* = 58.2; *p* < 0.00001).

**Table 1:** Effects of mating and receipt of sfAKH on insemination by a second male. Frequencies for females previously unmated, mated to wildtype (LVP) males, or mated to AKH-null males. Insemination by a second male was detected through the presence of red fluorescence (from dsRed males) in the female sperm storage organs. Numbers in table are the number of females with dsRed sperm out of the total tested. After their initial treatment, females were placed in cages with dsRed males for 24 hours.

| Treatment | Replicate 1 | Replicate 2 | Replicate 3 | Replicates Combined |
| --- | --- | --- | --- | --- |
| **Previously unmated** | 16 of 16 (100%) | 20 of 23 (87) | 11 of 11 (100) | 47 of 50 (94%) |
| **Previously mated to LVP** | 1 of 16 (6) | 0 of 15 (0) | 2 of 20 (10) | 3 of 51 (6) |
| **Previously mated to AKH-null** | 3 of 15 (20) | 5 of 25 (20) | 4 of 18 (22) | 12 of 58 (21) |

### 3.4 Females injected with MAG/SV from AKH-null males were more likely to be inseminated than females injected with MAG/SV from wildtype males

Since AKH-null males might differ from wildtype males in ways other than the presence of sfAKH, we repeated the previous experiment by comparing insemination patterns of females injected with MAG/SV homogenates from either wildtype or AKH-null males. In all three replicates of this experiment, 3-10% of females injected with MAG/SV from WT males, 7-29% of females injected with MAG/SV from AKH-null males, and 65-68% of females injected with saline were inseminated (Table 2). Insemination frequency did not differ between the replicates (*X^2^* = 1.30; *p* = 0.52), and, therefore, we combined the data from the three replicates for analysis. Females injected with MAG/SV of AKH-null males were significantly more likely to be inseminated than females injected with MAG/SV of WT males (*X^2^* = 5.3; *p* = 0.02). Females from both MAG/SV-injected treatments also were significantly less likely to be inseminated than saline-injected females (WT vs saline: *X^2^* = 70.0; *p* < 0.00001; AKH-null vs saline: *X^2^* = 43.9; *p* < 0.00001).

**Table 2:** Effects of injection with male accessory glands and seminal vesicles on female insemination. Frequencies for females injected with saline, wildtype (LVP) male MAG/SV, or AKH-null male MAG/SV. Insemination was detected through the presence of sperm in the female sperm storage organs. Numbers in table are the number of females with sperm out of the total tested.

| Treatment | Replicate 1 | Replicate 2 | Replicate 3 | Replicates Combined |
| --- | --- | --- | --- | --- |
| **Injected with saline** | 19 of 28 (68%) | 21 of 32 (66) | 15 of 23 (65) | 55 of 83 (66) |
| **Injected with LVP MAG/SV** | 2 of 34 (6) | 1 of 30 (3) | 3 of 30 (10) | 6 of 94 (6) |
| **Injected with AKH-null MAG/SV** | 10 of 34 (29) | 2 of 27 (7) | 3 of 32 (13) | 16 of 93 (17) |

### 3.5 Females injected with synthetic AKH peptide are less likely to be inseminated than females injected with a control peptide

To determine if AKH alone (i.e., in the absence of other seminal fluid molecules) impacts female insemination patterns, we compared insemination between females injected with AKH or a control peptide at two different concentrations. At 8pMol, there was no consistent trend for differences in the likelihood of insemination between females injected with the AKH or the control scrambled peptides (Table 3; *X^2^* = 0.14-0.84; *p* > 0.3). At 80pMol, in all three replicates with the LVP strain, 47-64% of females injected with AKH peptide and 66-100% of females injected with the control peptide were inseminated. Insemination frequency differed between replicates, so we did not conduct a combined analysis. This difference was only significant for one of the individual replicates (Replicate 2: *X^2^*= 11.0, *p* < 0.01). For the Thai strain, insemination frequency did not differ between replicates, so we combined replicates for analysis (*X^2^*= 3.0, *p* < 0.09). For the Thai strain, females injected with AKH were significantly less likely to be inseminated than females injected with the control peptide when the two replicates were combined (*X^2^* = 4.1; *p* = 0.04).

**Table 3:** Effects of injection of synthetic AKH peptide on female insemination. Frequencies for females injected with AKH or control peptide (8 or 80pMol) for wildtype (LVP or Thai) females. Insemination was detected through the presence of sperm in the female sperm storage organs. Numbers in table are the number of females with sperm out of the total tested.

| Treatment | Female Line Used |  |  |  |  |  |  |
| --- | --- | --- | --- | --- | --- | --- | --- |
|  | LVP<br>Replicate<br>1 | LVP<br>Replicate<br>2 | LVP<br>Replicate<br>3 | LVP<br>Replicates<br>Combined | Thai<br>Replicate<br>1 | Thai<br>Replicate<br>2 | Thai<br>Replicates<br>Combined |
| Injected<br>with 8pMol<br>AKH<br>peptide | 15 of 20<br>(75) | 14 of 24<br>(58) | 13 of 21<br>(62) | 42 of 65<br>(65) | N/A | N/A | N/A |
| Injected<br>with 8pMol<br>control<br>peptide | 11 of 18<br>(61) | 11 of 17<br>(65) | 18 of 24<br>(75) | 40 of 59<br>(68) | N/A | N/A | N/A |
| Injected<br>with<br>80pMol<br>AKH<br>peptide | 10 of 20<br>(50) | 16 of 25<br>(64) | 17 of 36<br>(47) | 43 of 81<br>(53) | 16 of 25<br>(64) | 21 of 26<br>(81) | 37 of 51<br>(73) |
| Injected<br>with<br>80pMol<br>control<br>peptide | 12 of 18<br>(67) | 25 of 25<br>(100) | 25 of 38<br>(66) | 62 of 81<br>(76) | 22 of 27<br>(81) | 29 of 31<br>(94) | 51 of 58<br>(88) |

### 3.6 Seminal fluid AKH does not impact likelihood or duration of remating

#### 3.6.1 The probability of remating did not differ based on a female’s previous receipt of sfAKH

We next investigated whether remating behavior differed between females previously mated to wildtype and AKH-null males. Out of all 182 females tested in this study, only 88 (48%) sustained genital contact for at least 6 seconds during the 4-minute trial and were, therefore, scored as mated (Table 4). In all three replicates of this experiment, 35-45% of females previously mated to WT males, 45-60% of females previously mated to AKH-null males, and 50-58% of previously unmated females were inseminated by a second male. Insemination frequency did not differ between the replicates (*X^2^* = 1.4; *p* = 0.49), and, therefore, we combined the data from the three replicates for analysis. There was no significant difference in the likelihood of remating across the three treatments (*X^2^* = 4.42; *p* = 0.11).

**Table 4:** Effects of mating and receipt of sfAKH on observed rematings with a second male. Frequencies for females previously unmated, mated to wildtype (LVP) males, or mated to AKH-null males. Rematings with a second male were determined through observations of individual females placed in cages of 20 males. Matings were scored as successful genital contact in the venter-to-venter position lasting at least six seconds. Trial duration was 4 minutes.

| Treatment | Replicate 1 | Replicate 2 | Replicate 3 | Replicates Combined |
| --- | --- | --- | --- | --- |
| Previously unmated | 12 of 21<br>(57%) | 10 of 20<br>(50) | 11 of 19<br>(58) | 33 of 60<br>(55) |
| Previously mated to LVP | 7 of 21<br>(33) | 7 of 20<br>(35) | 9 of 20<br>(45) | 23 of 61<br>(38) |
| Previously mated to AKH-null | 11 of 21<br>(52) | 9 of 20<br>(45) | 12 of 20<br>(60) | 32 of 61<br>(52) |

#### 3.6.2 The duration of remating did not differ based on a female’s previous receipt of sfAKH

There was no significant difference between replicates in mating duration (H = 0.71, *p* = 0.7), so we combined the replicates for analysis. Mating duration differed significantly across female treatment (H = 12.4, *p* = 0.002). Previously unmated females mated for significantly longer than females previously mated to WT (U = 580, *p* < 0.001) or AKH-null males (U = 738, *p* = 0.01). In all three replicates, females previously mated to AKH-null males had mating durations between those of unmated females or those previously mated to WT males (Figure 5). However, mating duration was not significantly different between females previously mated to WT or AKH-null males (U = 329.5, *p* = 0.40).

**Figure 5.**
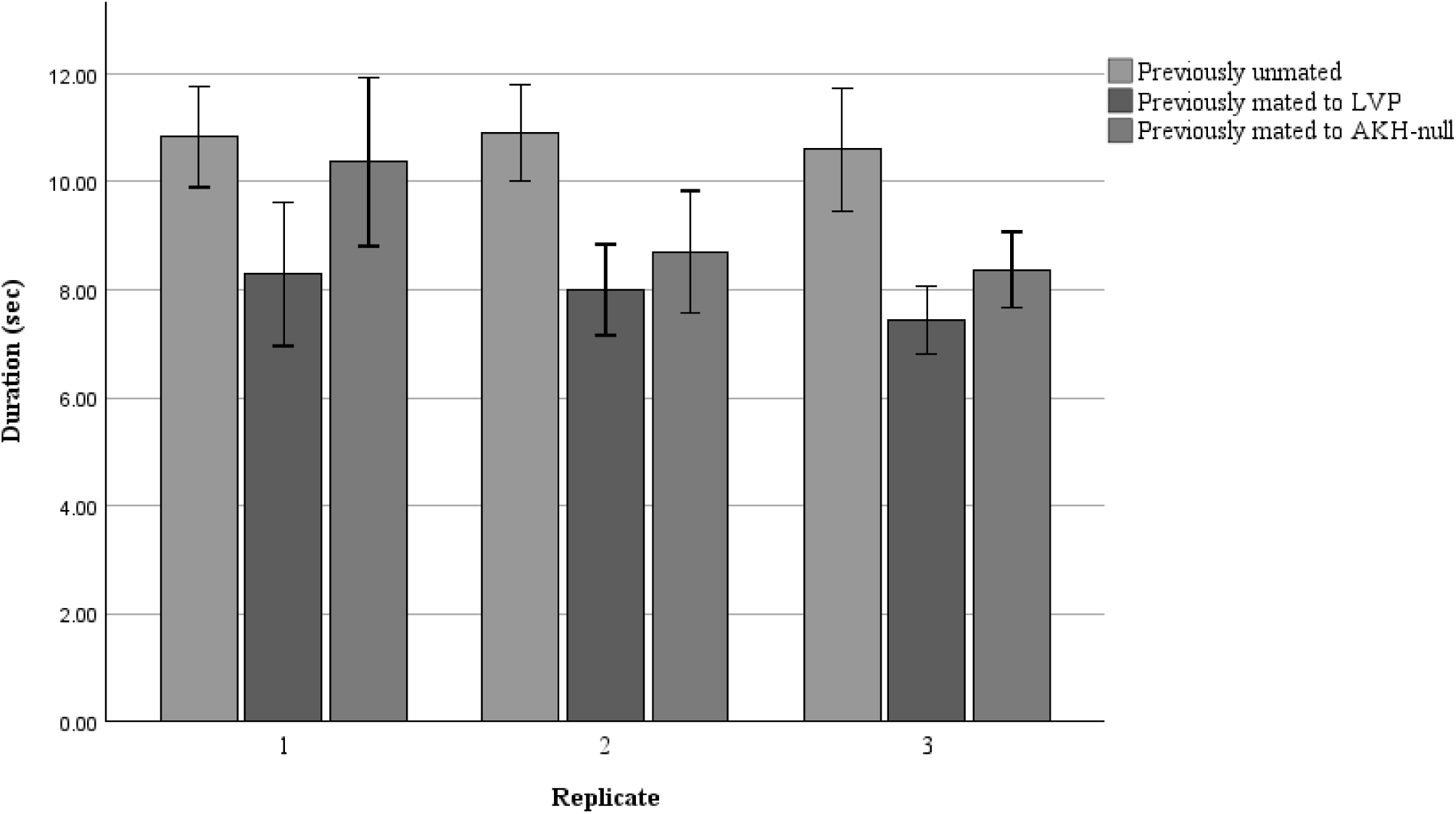
Mating duration (mean ± S.E.) of females in three treatments: previously unmated, previously mated to wildtype (LVP males), and female previously mated to males lacking the AKH peptide (AKH-null). N = 7-12 females per treatment per replicate; total N = 27-33 females per treatment.

## 4. Discussion

The AKH peptide is derived from a larger precursor protein that is comprised of a secretory signal sequence, then the AKH peptide, and finally a C-Terminus segment named the AKH precursor-related peptide (APRP; (Kaufmann et al., 2009; Marchal et al., 2018). Previously, we had discovered evidence for transfer of the APRP to female *Ae. albopictus* during mating (Boes et al., 2014). To date, no function has been discovered for the APRP (Clynen et al., 2004; De Loof et al., 2009; Galikova et al., 2015; Hatle & Spring, 1999). We had not detected the AKH peptide in our previous study either due to its absence or the inability of the technique we used to detect such a small peptide. Previous research in *Ae. aegypti* established that AKH mRNA is highly expressed in the abdomens of males and that the AKH receptor is expressed in both male and female reproductive tissues (Kaufmann et al., 2009; Oryan et al., 2018). In this study, we demonstrate the presence of the AKH mRNA and peptide in the male reproductive glands (SV/AG) and transfer of the AKH peptide from male *Ae. aegypti* to females during mating, thus establishing AKH as a novel *Ae. aegypti* seminal fluid molecule (SFM). This finding suggests an evolutionary transition in expression of this gene from exclusively in the *corpora cardiaca* to expression also in the male reproductive tract (Sirot, 2019). Such an evolutionary transition is not unprecedented in *Ae. aegypti*: males also produce juvenile hormone in their reproductive accessory glands (Borovsky et al., 1994). Together, these results suggest that sfAKH may have taken on a novel function in altering the post-mating responses of female *Ae. aegypti*.

We demonstrated an impact of AKH on female insemination patterns: females previously mated to AKH-null males were 3.5x more likely to be re-inseminated compared to females previously mated to WT males. Since male-derived AKH may impact other aspects of the male mating behavior, we tested the impact of injection of male reproductive glands from males with and without AKH and found similar results: unmated females injected with MAG/SVs of AKH-null males were 2.8x more likely to be inseminated compared to unmated females injected with the MAG/SVs of WT males. Finally, we tested the impact of injection with the AKH peptide on female insemination. Unmated females injected with a control peptide were 1.2-1.4x more likely to be inseminated compared to unmated females injected with the AKH peptide similarly reduced subsequent inseminations. Together, these results (i) demonstrate that long-term re-insemination refractoriness of female *Ae. aegypti* is induced, in part, by seminal fluid AKH (sfAKH) and (ii) confirm that full re-insemination refractoriness requires other contributions from the male MAG/SV (Fuchs et al., 1969; Lee & Klowden, 1999).

This system of female refractoriness being induced by a combination of SFMs of in *Ae. aegypti* shows similarities to *Drosophila melanogaster*, the most well-studied system for female post-mating responses (Avila et al., 2011; Wigby et al., 2020). In *D. melanogaster*, female insemination refractoriness results primarily from the receipt of a single SFM, sex peptide, which binds to sperm and is slowly released over the course of several days post-mating (Chapman et al., 2003; Liu & Kubli, 2003; Peng et al., 2005). However, for sex peptide to have its full effect, it requires sperm to bind to and other SFPs to bind it to sperm and facilitate its storage and release within the female (Peng et al., 2005; Ram & Wolfner, 2009; Sitnik et al., 2016). Further, there are other molecules transferred from males to females during mating that either decrease male mating attempts or increase female rejection behavior (Billeter & Wolfner, 2018; Saudan et al., 2002). Thus, *Ae. aegypti* and *D. melanogaster* are similar in requiring multiple SFMs to induce full female re-insemination refractoriness. Yet, these systems are different in that *D. melanogaster* refractoriness results from a single protein of major effect that is aided in its effect by other SFMs whereas *Ae. aegypti* refractoriness appears to result from partial effects of multiple SFMs that result in complete refractoriness when in combination (Fuchs et al., 1969; Lee & Klowden, 1999). This dependence on multiple proteins to induce refractoriness might be the legacy of a co-evolutionary arms race between male and female *Ae. aegypti* over control of re-insemination patterns (Sirot et al., 2015).

In addition to male contributions, re-insemination refractoriness of female *D. melanogaster* depends on many female contributions. These contributions include behavioral rejection of male mating attempts, the presence of sperm storage organs, and expression of the sex peptide receptor, neuromodulators, and other proteins in the nervous system (Chen et al., 2019; Häsemeyer et al., 2009; Rezával et al., 2012, 2014; Sirot et al., 2009; Sitnik et al., 2016; Yang et al., 2009; Yapici et al., 2008). Behavioral rejection of males also occurs in female *Ae. aegypti* in the form of kicking, abdominal twisting, wing-flicking, and not opening the vaginal plates (Aldersley & Cator, 2019; Craig, 1967; Cramer et al., 2023). Further, there is evidence that female *Ae. aegypti* may eject ejaculates after re-insemination attempts. During mating, males transfer sperm to the female’s bursa copulatrix. The sperm is subsequently stored in the spermathecae. Spielman et al. (1966) examined the internal and external genitalia of *Ae. aegypti* pairs that were flash frozen during rematings and found that semen was present in the females’ bursa during mating but absent after separation. Upon further examination, they found that semen was present between the females’ genital lips after the pair separated, suggesting that females are ejecting sperm during the separation process. The phenomenon of sperm expulsion should be further investigated in *Ae. aegypti*, as it is known to play an important role in modulating sperm dynamics in *D. melanogaster* (Manier et al., 2010). Other female contributions to re-insemination refractoriness in female *Ae. aegypti* are currently unknown.

Although we found that females previously mated to wildtype males were less likely to be re-inseminated than females previously mated to AKH-null males, we found no difference in the likelihood of remating between these two groups. In our study, we used a 6-second minimum duration of genital coupling as the criterion for a successful mating expected to result in insemination (Spielman et al., 1967). Our results suggest that, in the case of rematings, 6 seconds of apparent genital coupling does not necessarily result in insemination. This conclusion is supported by the results of the Spielman (1966) described above and the detailed observations of Houri-Zeevi et al. (Houri-Zeevi et al., 2025) in which they found tight interlocking of the male and female genitalia in matings involving previously unmated females but not in those involving previously mated females. This interlocking appears to be female-mediated in *Ae. aegypti*, as it requires females to elongate their genital tip. Female-mediation of the interlocking is also suggested by Craig (1967) who found that re-insemination was more likely in force-mated anesthetized females than in matings with awake females. Together with our findings, these results suggest that receipt of sfAKH may contribute to females rejecting male attempts to transfer semen during mating.

Our findings have implications for control strategies of *Ae. aegypti* that depend on the ability of males to inhibit female re-insemination, such as the sterile male technique. It may be possible to improve the ability of mass-reared males (e.g., for sterile insect or other control techniques) to inhibit female re-insemination by increasing the amount of sfAKH that the males produce. Although we have not yet tested this with sfAKH in *Ae. aegypti*, two lines of evidence suggest this approach could be effective. First, female insemination refractoriness depends on the amount of SFMs received: the higher the amount of SFM received, the lower likelihood of re-insemination (Helinski et al., 2012). Second, in *D. melanogaster*, artificial selection for larger male reproductive accessory glands resulted in higher production and transfer of sex peptide, an insemination inhibiting SFM (Wigby et al., 2009). Therefore, artificial selection for larger male reproductive accessory glands or genetic modification to increase sfAKH production could result in lab-reared males that are more effective in inhibiting female re-insemination. In sum, we have identified a novel seminal fluid protein in *Ae. aegypti* that our results indicate has a role in female insemination refractoriness. Further studies are needed to explore other female post-mating responses (e.g., feeding, digestion, flight behavior) that may be impacted by receipt of sfAKH.

## Acknowledgements

We thank Laura Harrington and Mariana Wolfner for providing feedback on an earlier version of this paper. We are grateful to James Becnel, Beth Lingenfelter, Jhony Mera, Nosherwan Mughal, and Tim Siegenthaler for administrative, logistical, and technical support. Research was supported by a U.S. National Institutes of Health grant R15AI140223. This project was part of the International Atomic Energy Agency Coordinated Research Project ‘D44005’. Graphical abstract created in Biorender.

